# Feasible closed-loop stimulation for neural latent dynamics control

**DOI:** 10.64898/2026.11.06.762216

**Authors:** Kyle A. Johnsen, Yijun Lai, Lavan Vivekanandasarma, Gordon J. Berman, Chethan Pandarinath, Christopher J. Rozell

## Abstract

Modern neuroscience benefits from powerful computational and experimental methods for studying neural populations, including latent dynamics models that predict behavior with few dimensions. But despite their explanatory utility, latent factors are seldom targeted explicitly with experimental perturbations, leaving causal relationships to behavior and inferred dynamical landscapes untested. A promising solution is closed-loop, multi-channel optogenetic control, which, when paired with model predictions, could steer neural trajectories in real time—a feat impossible for traditional, hand-designed stimulation. However, because closed-loop latent control has not been demonstrated *in vivo*, the hardware and software requirements remain unclear—particularly the necessary stimulation dimensionality and how limited resources such as implant volume and band-width should be allocated between stimulation and recording capabilities. Preliminary experiments would thus help guide ongoing technological development. As large-scale exploration of these design spaces *in vivo* is prohibitively expensive, we conducted a high-throughput simulation survey using biophysically grounded spiking neural networks. We found that low-dimensional actuation is sufficient: in the reference configuration, ~20% of the stimulation channels achieved ~85% of peak control performance, far fewer than the number of neurons. Furthermore, allocating channels roughly equally between stimulation and recording optimized behavioral control, whereas pairing many electrodes with a single light source—the prevailing practice in experimental labs—performed poorly. Even without heavily optimizing modeling or control, we obtained >60% tracking error reduction and >90% evoked behavior accuracy on average for optimal channel counts. These results held across extensive parameter variations, suggesting that closed-loop latent control is feasible with accessible channel counts and establishing design principles for future tool development and *in vivo* experiments.

## 1. Introduction

Recent advances in neural recording and stimulation technologies [1–5] have created unprecedented opportunities to measure and manipulate neural activity. These capabilities have catalyzed the latent dynamical systems paradigm, which explains high-dimensional population activity far better than intuitive, hand-crafted models [6–12]. Yet the latent state variables (“factors”) inferred by these models are rarely explicitly targeted via patterned stimulation. In systems neuroscience, some may think manipulating latent dynamics requires fast, single-neuron optogenetic resolution—driving heavy investment in two-photon technologies—alongside large-scale recording arrays to infer latent states.

Although *in vivo* experiments have achieved limited factor perturbations via open-loop or per-trial optimized stimulation [13–17], per-timestep feedback control offers multiple advantages (Fig. 1A). Namely full model-based optimization discovers per-timestep stimulation patterns that are difficult or impossible to design by hand, while real-time feedback compensates for disturbances and modeling errors. Optimal control also provides a quantitative framework for balancing tracking performance against other desiderata such as safety or remaining on the neural “manifold” [18, 19]. While closed-loop optogenetic control (CLOC), has been discussed for over a decade [20], it has historically targeted explicitly measurable variables—such as firing rate [21–23] or calcium fluorescence [24]—rather than population-level latent factors.

**FIGURE 1.**
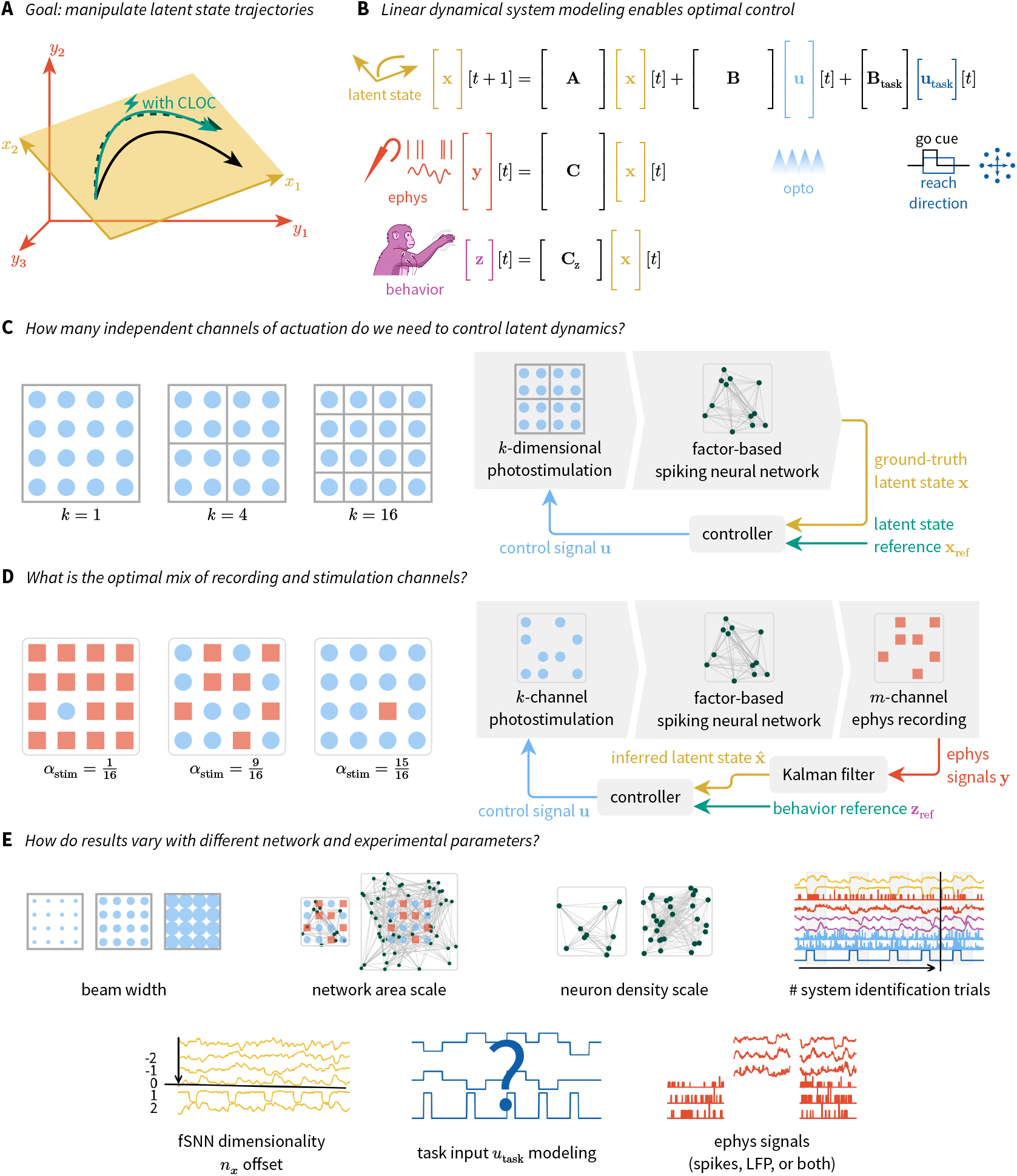
Overview of the objectives and design of this study. (A) The “manifold” of neural activity is represented by factors *x*_1_ and *x*_2_, forming a plane embedded in the higher-dimensional space of neural activity *y*_1_, *y*_2_, *y*_3_. We seek to steer factors **x** using closed-loop optogenetic control (CLOC). (B) Linear dynamical system modeling, where the latent state is driven by optogenetic and task inputs and maps to neural and behavioral observations, serves as the basis for our control experiments. for simplicity here, a 32 × 32 grid was used in experiments. Right: A control diagram for the same (C) Our first experiment probes how individuated stimulation must be to control latent dynamics. Left: We use the same grid of light sources across conditions, traversing between the two extremes of treating all as a single actuator and treating each as a separate actuator. While a 4× 4 grid is shown for simplicity here, a 32 × 32 grid was used in experiments. Right: A control diagram for the same experiments, where the controller is given ground-truth factors from the network and a reference trajectory defined in latent space. (D) The second experiment quantifies optimal allocation of a fixed channel budget between recording and stimulation for optimal control. Left: Experimental settings range from one channel used for stimulation and the rest for recording to the opposite extreme. While a 4 × 4 grid is shown for simplicity here, a 10×10 grid was used in experiments. Here and throughout, blue circles represent light sources and red squares represent electrode contacts. Right: A control diagram for these experiments, differing from *C* in that latent states are estimated from measurements and the reference trajectory is defined in terms of behavior. (E) We repeat experiments for each of the parameters represented here to assess the robustness of our findings and approach. The electrophysiology signals parameter only applies in the stimulation–recording balance experiments.

To create a framework for successful closed-loop latent factor control, critical technical unknowns must be solved: First, is low-dimensional actuation sufficient to navigate similarly low-dimensional latent manifolds? While high-fidelity, per-neuron control would be necessary to stay on the full, high-dimensional neural manifold [19], whether such fine-grained actuation is required to traverse a lower-dimensional “manifold of interest” defined by behavior-correlated factors remains an open question [18]. Second, as recording channel counts surge [3], how should an experimental interface allocate a fixed channel budget between stimulation and recording for maximally informative experiments? Access to the brain is limited by implant footprint, wiring capacity, and thermal constraints, making efficient use of stimulation and recording resources important for interface design. These constraints differ between modalities but can overlap, coupling the resources available for sensing and actuation. Although the optimal allocation depends on the specific application and hardware, even the general principles governing this tradeoff remain an open question.

Resolving these questions would thus be valuable, but *in vivo* experiments in trained animals and prototype hardware are expensive and pose a substantial engineering challenge— especially given the need for large parameter sweeps to systematically explore the space of problems. Therefore, we sought to answer these questions instead by conducting a high-throughput survey of end-to-end, *in silico* CLOC experiments using the Cleo simulator [25]. Cleo incorporates models of multi-electrode recording, optogenetic stimulation, and closed-loop control into spiking networks, providing a level of realism absent from abstract control studies with non-spatial, non-biophysical inputs and outputs [26]. We embedded these interfaces in factor-based spiking neural networks (fSNNs) [27], which define latent factors as explicit readouts of network activity, interacting through low-rank connectivity. These networks model a generic cortical patch without distinct cell types, laminar structure, or cell-type-specific opsin expression, isolating the role of spatial actuation granularity. fSNNs were trained to produce low-dimensional latent dynamics as they follow a center-out reaching task, with behavioral readouts approximating macaque electromyography (EMG) data. Simulations modeled the entire experimental pipeline: spatial interface placement, random stimulation to produce system identification data, linear dynamical system (LDS) fitting (Fig. 1B), and closed-loop model predictive control (MPC) to guide latent and behavioral trajectories. During control experiments, the controller attempted to track reach trajectories in randomly sampled directions while the network lacked a directional cue.

With this pipeline, we conducted two main sets of experiments. First, we varied the number of independently controlled light sources, ranging from a single spatially uniform channel to one channel per neuron, to assess the stimulation dimensionality required to control latent factors (Fig. 1C). In the reference configuration, we found that ~20% of the stimulation channels achieved ~85% of maximum control performance on average. This suggests that fine-grained, single-neuron stimulation is unnecessary for steering low-dimensional latent dynamics, challenging prevailing technological priorities. Second, we evaluated the allocation of a fixed 100-channel budget between stimulation and recording for behavioral control (Fig. 1D), assigning equal nominal channel costs to probe fundamental sensing–actuation tradeoffs. This equal weighting provides a starting point for future application-specific studies, in which stimulation and recording costs would be weighted according to their respective hardware constraints. Under this simplified budget, we found that an approximately equal balance optimized performance. In contrast, the low-actuation configuration ubiquitous in systems neuroscience—many recording electrodes paired with a single optical fiber—provided little improvement over baseline. These findings remained robust across thousands of simulated trials sweeping neuronal density, optical beam width, target factor count, and the amount of training data (Fig. 1E). Even with standard linear control and minimal tuning, optimal channel counts achieved ~66% tracking error reduction and ~94% reach direction classification accuracy in the reference configuration. Together, these results provide evidence for the feasibility of low-dimensional latent factor control and help inform experiment and tool design, laying groundwork for more powerful causal study of latent dynamics hypotheses.

## 2. Methods

### 2.1. Experimental pipeline overview

To investigate the requirements for closed-loop optogenetic control of latent dynamics, we simulated end-to-end experiments under two primary experimental paradigms:

a. **Stimulation dimensionality experiments:** We evaluated how many independent actuation channels are needed to steer latent factors (Fig. 1C). A fixed grid of 1,024 light sources (32 × 32, 4 mm×4mm) was grouped into *k* ∈ [1, 1024] independently controlled channels, ranging from uniform illumination (*k*= 1) to individual source control (*k*= 1024).
b. **Stimulation–recording balance experiments:** We evaluated how to optimally partition a fixed budget of 100 channels (10 ×10 grid), motivated by implant volume and bandwidth constraints, between *k* stimulation light sources and *m* recording electrode contacts such that *k* + *m* =100 (Fig. 1D). Here, the controller lacks ground-truth factors and must infer latent states from electrode recordings to control behavioral outputs.

For each condition, simulations followed a five-stage experimental workflow (Fig. 2): first, generating and training a factor-based spiking neural network (fSNN) to reproduce macaque reach kinematics via low-dimensional latent factors (Fig. 2A); second, interfacing the network with Cleo to model spatial optogenetic stimulation and electrophysiological recording (Fig. 2B); third, delivering random photostimulation during system identification trials to collect input–output data (Fig. 2C); fourth, fitting a linear dynamical system (LDS) model to capture stimulus-evoked latent dynamics; and fifth, simulating closed-loop model predictive control (MPC) or open-loop (OL) stimulation to compensate for removed task cues and guide the network along target trajectories (Fig. 2D). The following subsections describe each stage of this pipeline in detail.

**FIGURE 2.**
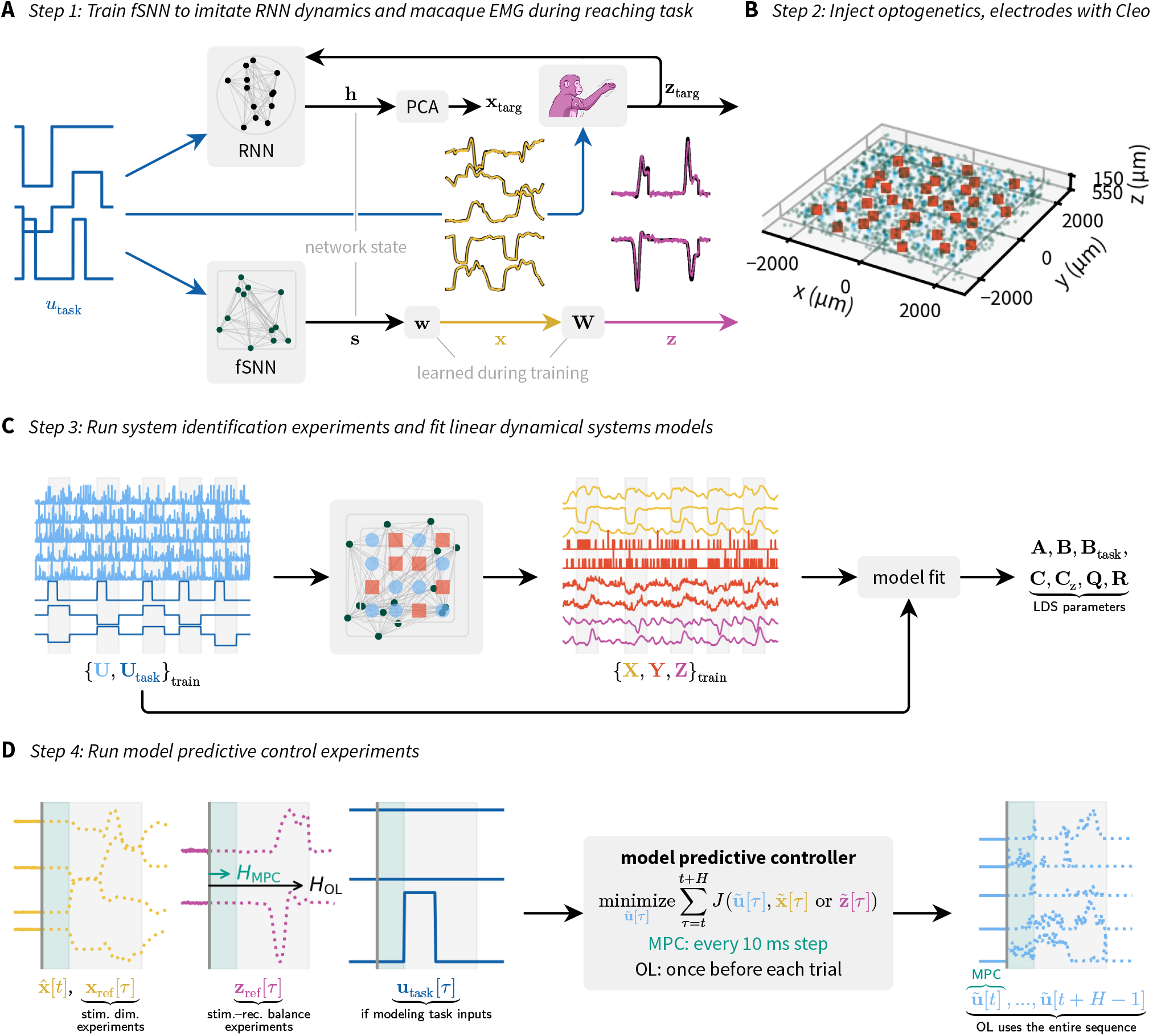
An end-to-end pipeline for simulating closed-loop optogenetic control of latent dynamics. (A) A factor-based spiking neural network (fSNN) is trained to produce low-dimensional dynamics and EMG signals during a reaching task. Task inputs (go cue and target direction) and EMG from macaque experiments are fed into a recurrent neural network with random weights (top). The top principal components of the hidden state trajectories are used as factor targets **x**_targ_ for the fSNN, while the EMG serves as the behavioral target **z**_targ_. The same task inputs are also fed into the fSNN (bottom), while the neuron-factor and factor-behavior weight matrices **w** and **W** are updated recursively to match the targets. (B) The trained fSNN is then interfaced with simulated optogenetics and, in stimulation–recording balance experiments, electrophysiology. Light sources and electrode contacts are arranged in a grid pattern, across a square slab of neurons with random coordinates. (C) Random photostimulation is delivered along with standard reaching task inputs during 100 trials of system identification experiments, and the resulting data is used to fit linear dynamical system (LDS) models. (D) 100 trials of control experiments are then simulated, where a model predictive controller receives the current state estimate, the reference trajectory over the control horizon, and optionally the task inputs. The reach direction task inputs are removed during control experiments, such that the controller effectively attempts to compensate as it follows the reference trajectory. Informed by LDS parameters, the controller optimizes the future control trajectory to minimize tracking error and input power, subject to constraints on the stimulation. The same optimization problem governs feedback control (MPC) and open-loop control (OL): MPC re-solves every 10 ms with a 300 ms (30-step) rec5eding horizon, whereas OL solves once per trial over the entire 1.11 s (111-step) duration.

### 2.2. Factor-based spiking neural network models

To endow our models with meaningful low-dimensional dynamics, we implemented factor-based spiking neural networks (fSNNs) [27]. An fSNN’s low-rank recurrent weights are trained via recursive least squares to reproduce low-dimensional target factors, defined as weighted sums of filtered spike trains across all neurons. We simulated a center-out reaching task based on experiments where non-human primates were shown one of eight targets and initiated a reach after being shown a go cue [28, 29]. Condition-averaged electromyography (EMG) traces from the lateral biceps and posterior deltoid served as behavioral targets (**z**_targ_) for learning an affine readout from factors to outputs. Factor targets (**x**_targ_) were the first *n*_*x*_ principal components capturing 99% of the variance in the hidden state trajectory of a rate-based recurrent neural network (RNN). The RNN received direction cues, the go cue, and EMG outputs as input, and principal components were computed on 100 trials of preliminary data (Fig. 2A). The *n*_*x*_ offset exploratory parameter adjusted the number of factors trained and controlled; because additional factors represent lower-variance components, *n*_*x*_ offset serves as a weak proxy for network complexity and control difficulty (Fig. 1E).

Neurons, without distinct cell types or laminar structure, were randomly distributed within a 400 µm-thick volume at a base density of 125 neurons/mm^3^, producing 1,024 neurons in the reference configuration—equal to the number of light sources. Biologically realistic densities (e.g., ~65,000/mm^3^ [30]) were computationally intractable given dense network connectivity. The density and network area scale exploratory parameters scaled neuronal density relative to the base and network footprint area relative to the 4 mm ×4 mm interface array (Fig. 1E).

### 2.3. Simulated experimental interface

After training, the experimental interface was then defined with Cleo, a closed-loop control, electrophysiology, and optogenetics simulator [25] for Brian 2 [31] spiking network models (see Fig. 2B). Cleo facilitates experiment simulation by taking a user-provided network and injecting models of light propagation, opsins, spike detection, local field potential recording, calcium imaging, closed-loop latency, etc., sparing the user from implementing them ad hoc. In stimulation dimensionality experiments, the 32×32 light grid was grouped into *k* contiguous clusters using constrained Voronoi partitioning (Sec. A.1). This preserved total light emission across channel counts *k*, varying only spatial granularity (Fig. 1C). In stimulation–recording balance experiments, the 10 ×10 grid was partitioned into *k* light sources and *m* electrodes using an iterative repulsion algorithm that maximized distances between channels of the same modality (see Fig. 1D, Sec. A.1 for details). Light sources followed an optical fiber propagation model for 473 nm (blue) light [32]. Photostimulation delivered a current proportional to local irradiance for each neuron, assuming uniform opsin expression, scaled so that average task input (mean [|*w*_in|]_) matched a 5 mW/ mm^2^ light input 500 µm below the fiber tip. Stimulation was capped at 75 mW /mm^2^, as may be done *in vivo* to avoid tissue damage [33].

Electrodes recorded multi-unit activity (MUA), LFP, or both. MUA detection followed Cleo defaults, with three adjustments: a (longer) 200 µm noise floor radius, a (lower) 3σ_noise_ spike detection threshold, and a (higher) 0.01 recall cutoff. We label these threshold-crossing MUA recordings “spikes” in figures. The first two compensated for network sparsity while the third increased computational efficiency. LFP was simulated as a distance-weighted sum of total synaptic currents across neurons:

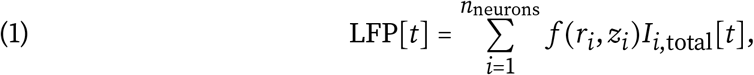

where the weight *f* is a function of the lateral and axial distance *r* and *z* from the neuron to the electrode.

### 2.4. System identification experiments

We then simulated system identification experiments to obtain data for training dynamical systems models of the network (see Fig. 2C). We varied *n*_trials sys ID_ to measure how the amount of training data affects the success of control experiments. Trials followed the same pattern of the original experiment—target presentation, movement following a go cue, and an intertrial rest period—but with random optogenetic stimulation throughout. Stimulation was sampled independently per channel, following a right-censored log-normal distribution

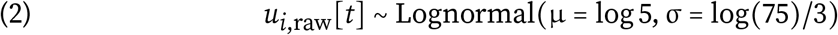

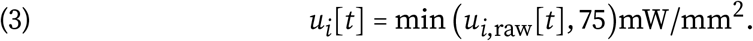

Photostimulation was updated and measurements were recorded every 10 ms, in contrast with the base 1 ms time step of the underlying fSNN simulation.

### 2.5. Dynamical system model fitting

Linear dynamical system (LDS) models were then fit to the training data for later use in optimal control experiments. Training data was normalized to zero mean and unit variance per channel before fitting. LDS models took the form

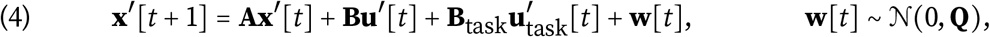

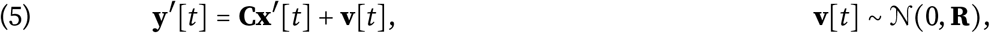

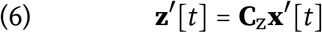

where ′ indicates normalized LDS variables, as opposed to the original quantities used in the fSNN, 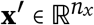 are the latent factors, **u**′∈ **R**^*k*^, 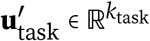 are the stimulation and task inputs, 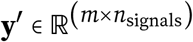 are the recorded measurements (MUA and/or LFP), and **z**^′^ ∈ **R**^3^ are the behavioral outputs (two EMG channels and one stabilizing signal). **A, B, C**, and **C**_z_ are the system, input, measurement, and behavioral output matrices, respectively, while **Q** and **R** are the process and measurement noise covariances. Here we varied the task input modeling parameter, which determined whether task input channels were included in **u**′ during fitting and later state inference and control. When included, present and future task inputs were known perfectly.

In stimulation dimensionality experiments, **A, B, C**, and **C**_z_ were fit via least squares using ground-truth network factors, with **Q** and **R** estimated from residuals. Using ground truth eliminated suboptimal model fitting, directly isolating the impact of actuation di-mensionality. In stimulation–recording balance experiments, where latent states must be inferred from recordings, we fit models using BRAID [34]. BRAID explicitly identifies latent factors predictive of behavior—useful in a realistic setting where the most interesting factors to manipulate would be those correlating with behavior. We used standard linear BRAID with one-step-ahead prediction loss and no behaviorally irrelevant factor estimation.

### 2.6. Control experiments

Fitted LDS models were then used to perform 100 trials of optimal control experiments. We simulated reaching trials but removed the directional task cue, requiring the controller to guide the reach via optogenetics alone. In stimulation dimensionality experiments, where ground truth was available, the reference trajectory was defined in latent space: **x**_ref_ [*t*]*=***x**_targ [_*t*]. In stimulation–recording balance experiments, where inferred factors were not directly comparable to ground-truth factors, we instead defined the reference in terms of behavior, multiplying the rate network target factors by the fSNN’s output weights: **z**_ref_ [*t*] *=* **Wx**_targ_ [*t*]. Note that **x**_targ_ (RNN-derived target factors), **x** (the fSNN’s ground-truth factors), and 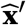 (factors inferred from recordings) are distinct quantities despite the shared *x* notation. Controlling behavior directly captures the trade-off between state inference accuracy (from recording channels) and state controllability (from stimulation channels).

Control was implemented via a model predictive control (MPC) framework, using quadratic programming to solve for the optimal sequence of future control inputs over a finite horizon at each control step (see Fig. 2D). The problem is formulated as

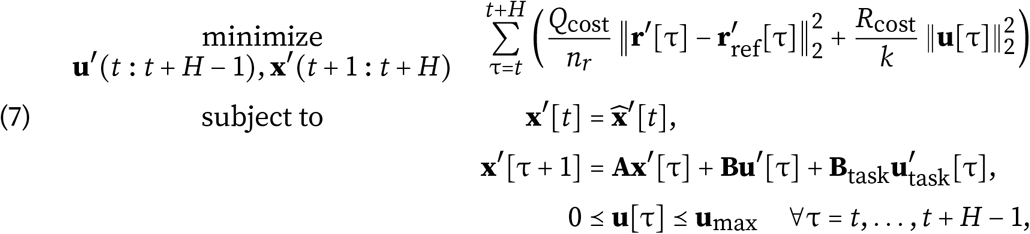

minimizing normalized and weighted tracking error and input magnitude, subject to the fitted LDS dynamics and constraints on **u [***t*] (non-negativity and maximum intensity of 75 mW mm^2^ per channel, where ≤ denotes elementwise inequality and **u[***t*] is unnormalized). **r** is the reference variable (**x** in stimulation dimensionality experiments or **z** in stimulation–recording balance experiments), *H* is the control horizon length, **r**_ref_ [*t*] is the reference variable at time *t*, and *Q*_cost_ and *R*_cost_ are the tracking error and input cost scalars, respectively.

Before simulation, OSQP [35], Clarabel [36], and SCS [37] solvers were benchmarked and ranked by median solve time, falling back to subsequent solvers upon failure. Solve time limits (including a minimum 1 ms baseline and solver overhead) were set to 20 ms for MPC and 2 s for open-loop (OL) control. If the budget was exceeded, the solver returned the current suboptimal iteration (observed only occasionally during high-*k* MPC). The controller was active from 300 ms before target presentation through trial end. For open-loop control, the optimization was solved once per trial. For closed-loop MPC, it was re-solved every 10 ms using updated state estimates, applying only the first step’s control input (Fig. 2D). State estimates 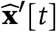 were obtained from fSNN ground truth in stimulation dimensionality experiments and Kalman filtering [38] in stimulation–recording balance experiments (Fig. 1C,D).

### 2.7. Exploration of parameter space

The entire suite of experiments was performed for each value of *k* (the number of stimulation channels), each of eight different random seeds, and each control type (MPC with/without latency, and open-loop). This constitutes the main result set, with a number of exploratory experimental and network parameters (Fig. 1E) set to a reference configuration. Then, starting from the reference condition, each parameter was swept individually, exploring the parameter space in a “one-factor-at-a-time” (OFAT) pattern. Table S1 lists these exploratory parameters, their reference values, the range of values swept, as well as many other parameters that were held constant.

### 2.8. Data analysis

Control performance was evaluated using normalized tracking quality *Q*_track_(**r**) ∈ (−∞, 1]:

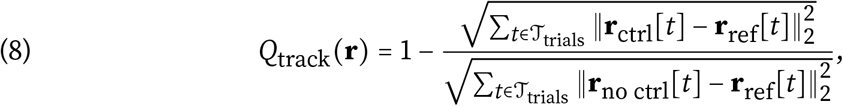

where **r**_ctrl_ and **r**_no ctrl_ represent trajectories with active control and zero stimulation, respectively. Here, *Q* 1 indicates perfect tracking, *Q* 0 denotes performance equivalent to zero stimulation, and *Q* 0 indicates tracking worse than baseline. We evaluated 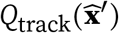 for stimulation dimensionality experiments and *Q*_track_ **z** for stimulation–recording balance experiments.

We also quantified control performance for stimulation–recording balance experiments in terms of the evoked behavior via the prediction accuracy of a *k*-nearest neighbor classifier. The classifier was “trained” with the reference trajectories **z [***t*] as inputs and the true conditions (reach directions) from control experiment data as the labels and “tested” with the evoked **z**[*t*] trajectory as the inputs. Classifier success thus meant that the evoked trajectory was closer to reaches of the intended direction than other directions, among the *k* =3 nearest neighbors.

Key metrics for each experiment set are derived from *Q*_track_. To quantify to what extent low-dimensional actuation can control latent dynamics, we compute *k*_knee_, the number of stimulation channels *k* of the “knee” point in the curve. This is defined as the point of maximum curvature, providing an objective measure of diminishing returns with increasing stimulation granularity. We obtained *k*_knee_ via a modified Kneedle algorithm [39], which, unlike the original algorithm that takes the first or the last candidate knee point, takes the candidate with the highest difference measure (maximum curvature). To quantify optimal stimulation–recording balance, we simply report the best-performing proportion of channels allocated to stimulation 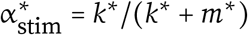, where *k*^*^, *m*^*^ = argmax_*k,m*_ *Q*_track(_ *k, m*) are the number of stimulation and recording channels, respectively, that achieve the best control performance.

To quantify the extent to which metrics are affected by each parameter, we compute the conditional mutual information (CMI) between the parameter and the metric of interest, conditioning on the random seed and, in the case of control performance, the number of stimulation channels *k*. We choose CMI because it captures nonparametric, nonmonotonic relationships. We estimate CMI *I* (*A*; *B*| *C*)with a plug-in estimator, taking the average over local estimates:

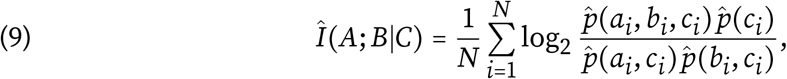

where 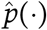 represents the density obtained via kernel density estimation (KDE) and *A, B, C* represent the exploratory parameter, the metric of interest, and the conditioning variable(s), respectively. For details on the KDE implementation, see Sec. A.2.

To estimate statistical significance of estimated CMI values, we performed a permutation test for each exploratory parameter. Within each seed, *k* group, we shuffled the order of the metric (*B* in Eq. 9) to break any dependence relationship and computed the CMI 1,000 times. We computed the *p*-value for each parameter as the proportion of the shuffled CMI values that were equal to or higher than the point estimate for the unshuffled data. To correct for multiple comparisons, we computed Benjamini-Hochberg false discovery rate-adjusted *q*-values [40] for each of the four result sets (2 experiment paradigms ×2 metrics).

### 2.9. Computational tools

Experiments were performed on a high-performance computing cluster whose nodes are equipped with 64 GB RAM, dual Intel Xeon Gold 6226 CPUs @ 2.7 GHz (24 cores/node), DDR4-2933 MHz DRAM, and Infiniband 100HDR interconnect. The cluster operating system was Red Hat Enterprise Linux 9.6 and Slurm [41] was used for job scheduling. The multi-step pipeline was orchestrated with Snakemake [42], which efficiently handles dependencies between steps and parallel cluster job submission. Spiking network simulations were implemented with Brian 2 [31] and electrophysiology and optogenetics simulations with Cleo [25] v0.18.3. Network training was accelerated with the Brian2CUDA GPU backend [43] on NVIDIA V100, A100, and Quadro RTX 6000 GPUs. Code for the entire pipeline and analysis will be made publicly available at https://github.com/siplab-gt/control-of-latent-dynamics upon publication, including an environment snapshot with exact dependency versions to ensure reproducibility.

## 3. Results

In both sets of experiments, closed-loop optogenetics successfully steered network dynamics along target trajectories. Representative trajectories (Fig. 3) exhibit high trial-to-trial consistency per condition, reflecting stable network dynamics and control.

**FIGURE 3.**
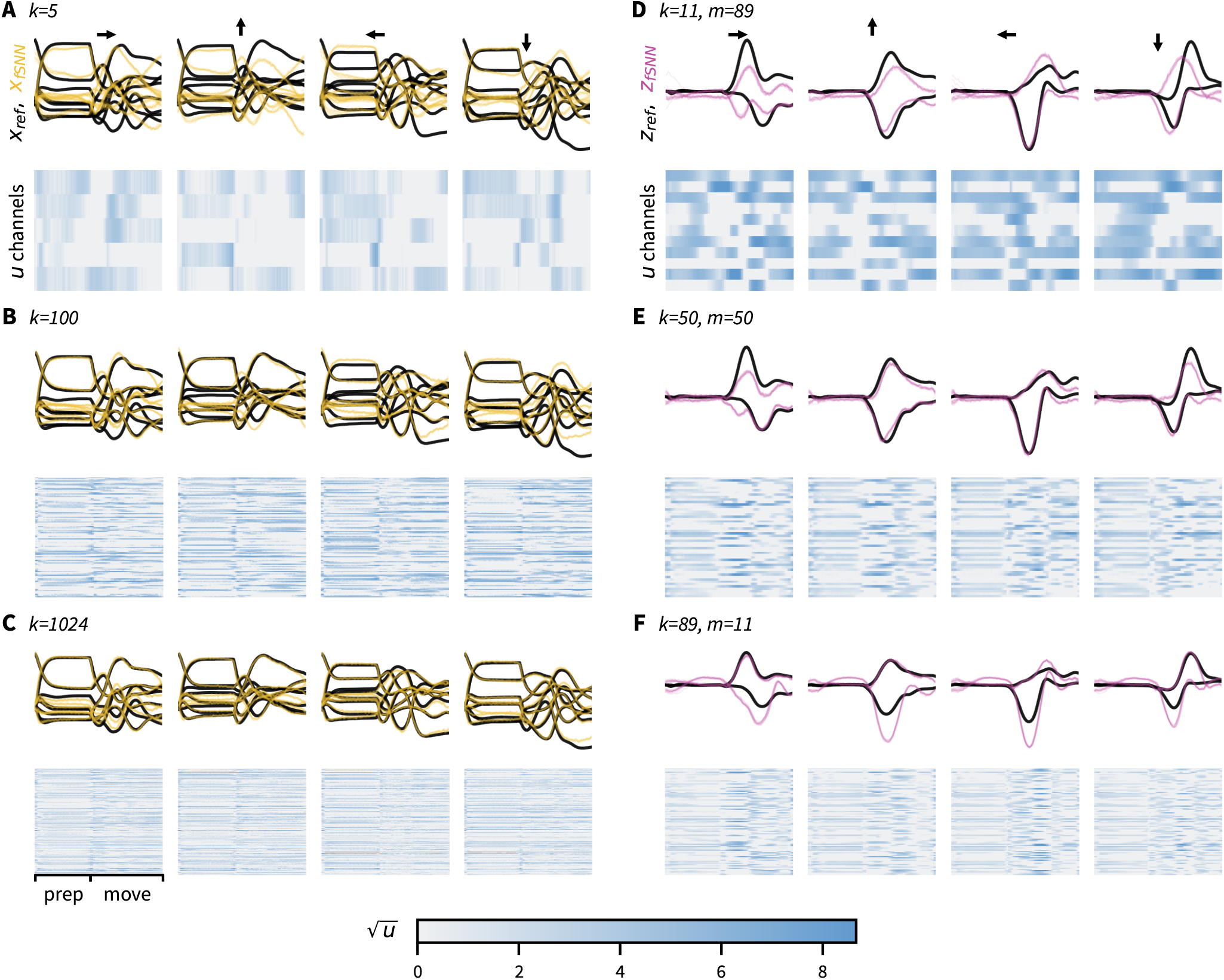
Representative example control trajectories under the reference configuration (Table S1), showing high trial-to-trial consistency. (A-C) Trajectories for low-, medium-, and high-dimensional stimulation (*k* 5, 100, 1024, respectively) in stimulation dimensionality experiments. Four reach directions—left, up, right, down—are shown, one in each column. Top: latent state trajectories, with the average reference in black (s.e.m. too small to visualize). All trials are overlaid for each condition in thin, colored lines, with factors on the y axis. Bottom: condition-averaged light stimulus patterns are shown in heatmaps with channels on the y axis. The x axis in both represents time within each trial, including the preparation and movement periods. (D-F) Trajectories for low-, medium-, and high-dimensional stimulation ratio (α 0.11, 0.5, 0.89, respectively) in stimulation–recording balance experiments. The total channel count is *k* + *m =*100. Plots and heatmaps are as in *A-C*, but with behavior instead of factors. Data shown from random seed 16320829.

### 3.1. Adequacy of low-dimensional actuation for controlling latent dynamics

For stimulation dimensionality experiments, we ran the complete pipeline described above for channel counts *k* ∈ [1, 1024], repeated for each of 8 different random seeds. We found a recurring pattern of diminishing returns (concave curves) when plotting performance vs. channel count. In the reference configuration, the curve reached its knee point (point of maximum curvature) at *k*_knee =_200 (the same tested grid point in all eight seeds) independently controlled light-source groups^1,2^ (Fig. 4A). This knee point was lower than the neuron count, though larger than the latent dimensionality (*n*_*x*_< *k*_knee_< *n*_neurons_). Crucially, these knee-point channel counts achieved 85.0 ±1.91% of peak tracking capacity 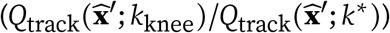.

**FIGURE 4.**
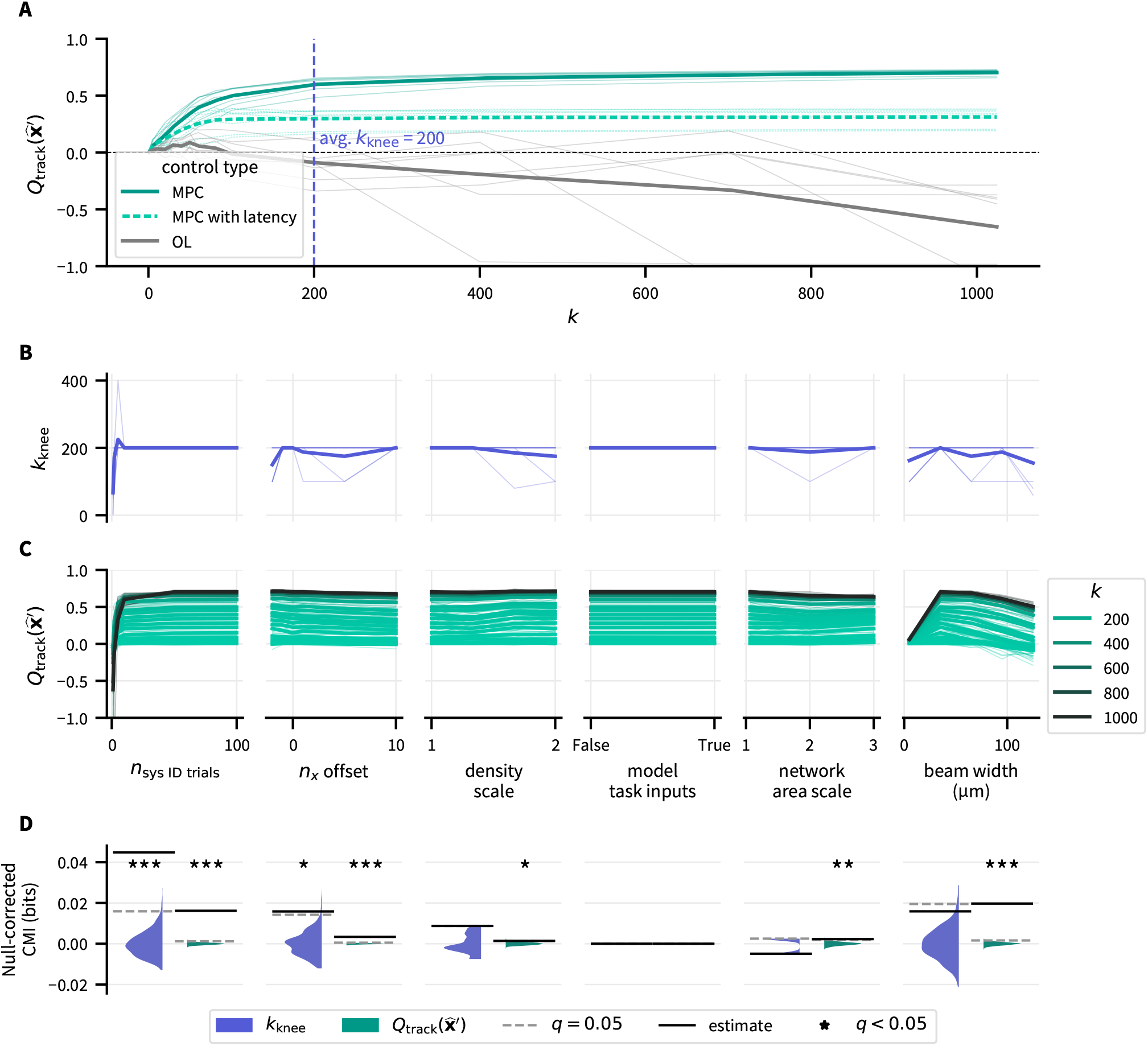
Diminishing returns are observed at low channel counts across a wide range of conditions. (A) The average knee point of the performance/channel count curve reaches relatively high performance with a low number of channels. The curve for control with latency plateaus near *Q*≈ 0.30 after *k*= 100 while no-latency control keeps rising, demonstrating that practical compute constraints could nullify the small advantage provided by higher channel counts. Open-loop (OL) control performed near baseline at low channel counts and below baseline at high channel counts, suggesting that multi-output latent dynamics trajectory control requires intra-trial feedback. Here and throughout, thin lines indicate individual seeds and thick lines indicate the mean across seeds. (B) The knee point of the curve is relatively consistent across exploratory parameters, with the factor count *n*_*x*_ offset and system identification trial count showing significant dependence. (C) Control performance depended significantly on all exploratory parameters except task input modeling. (D) Permutation tests, where metric values are shuffled with respect to the parameter values per seed (and channel count for control performance) show statistically significant non-independent relationships. Histograms show CMI estimates for each of 1,000 permutations, solid lines show the point estimate of the unshuffled data, and dashed lines show the *q* =0.05 significance threshold. *, **, *** indicate *q* < 0.05, 0.01, 0.001, respectively.

This diminishing-returns profile remained consistent across exploratory parameter sweeps, with two statistically significant exceptions (Fig. 4D).

- The knee point *k*_knee_ depended significantly on the factor count *n*_*x*_ offset, without a consistent monotonic trend.
- System identification trial count also showed a significant dependence, with low trial counts yielding smaller *k*_knee_ estimates and high variance in the low-trial regime (Fig. 4B).

While *k*_knee_ reflects channel efficiency, overall control performance *Q* measures trajectory tracking accuracy. Analyzing *Q*, we observe two notable patterns. First, when simulating compute latency, *Q* plateaued near *k*= 100 in the reference configuration, demonstrating how computational delays can limit the advantage of higher channel counts. Second, performance varied across exploratory parameter sweeps. Notable dips occurred for low system identification trial counts, high *n*_*x*_ offset, and extreme beam widths. Permutation tests identified statistically significant conditional mutual information (CMI) dependencies for all exploratory parameters except task input modeling (see Fig. 4C,D, Fig. S1). Poor control is expected for little training data, since the resulting inaccurate model would inform the controller poorly. Extreme beam sizes degraded tracking: narrow beams recruited too few neurons to steer dynamics, while wide beams’ recruitment of too many neurons harmed factor selectivity. Detailed CMI statistics are provided in Table 1.

**TABLE 1.**
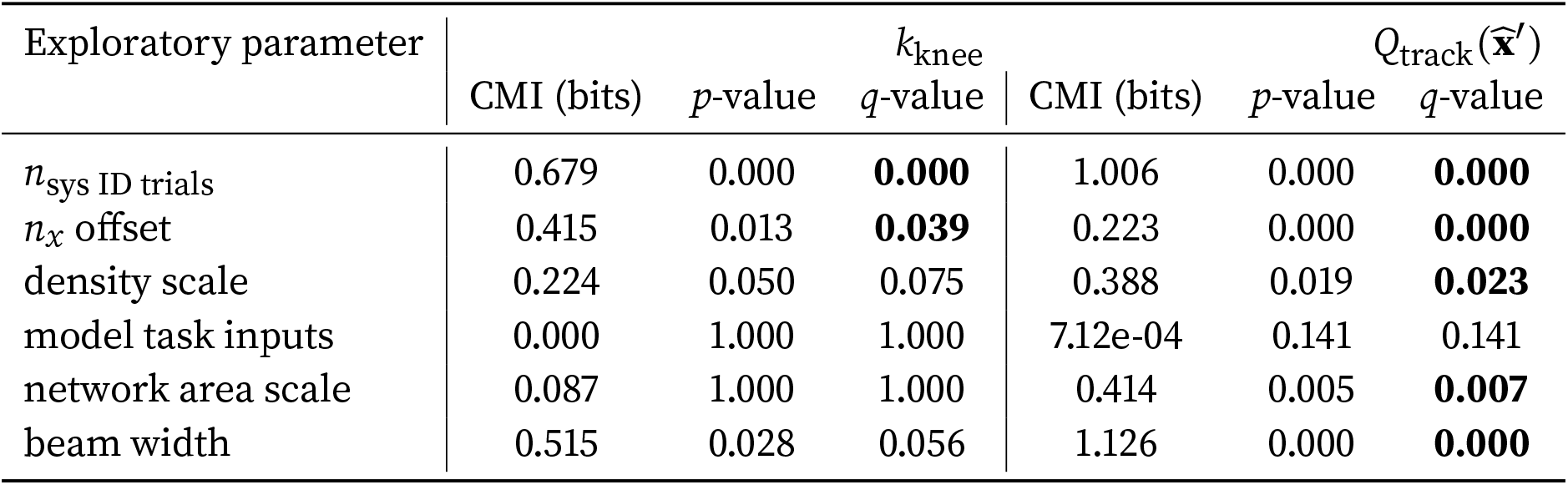
Statistical analysis results for stimulation dimensionality experiments, relating *k*_knee_ and 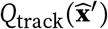 to exploratory parameters.

| Exploratory parameter | *k*<sub>knee</sub> |  |  | *Q*<sub>track</sub>(**x̂**′) |  |  |
|---|---|---|---|---|---|---|
|  | CMI (bits) | *p*-value | *q*-value | CMI (bits) | *p*-value | *q*-value |
| *n*<sub>sys ID trials</sub> | 0.679 | 0.000 | **0.000** | 1.006 | 0.000 | **0.000** |
| *n*<sub>*x*</sub> offset | 0.415 | 0.013 | **0.039** | 0.223 | 0.000 | **0.000** |
| density scale | 0.224 | 0.050 | 0.075 | 0.388 | 0.019 | **0.023** |
| model task inputs | 0.000 | 1.000 | 1.000 | 7.12e-04 | 0.141 | 0.141 |
| network area scale | 0.087 | 1.000 | 1.000 | 0.414 | 0.005 | **0.007** |
| beam width | 0.515 | 0.028 | 0.056 | 1.126 | 0.000 | **0.000** |

### 3.2. Optimal balance of stimulation and recording for controlling latent dynamics

In stimulation–recording balance experiments, we swept stimulation channels *k* ∈[1, 99] (and recording channels *m=* 100 −*k*) across 8 seeds for both reference and exploratory configurations. In the reference configuration, optimal behavioral control required a balanced channel allocation with a slight stimulation bias: optimal stimulation channel proportion 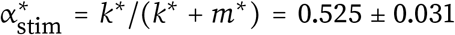 (Fig. 5A). And, while we cannot judge whether control performance is sufficient without a specific scientific application, the 66.2 ±2.28% tracking error reduction/93.6 2.63% reach direction classification accuracy^3^ achieved at optimal channel counts is conceivably adequate for many scenarios.

**FIGURE 5.**
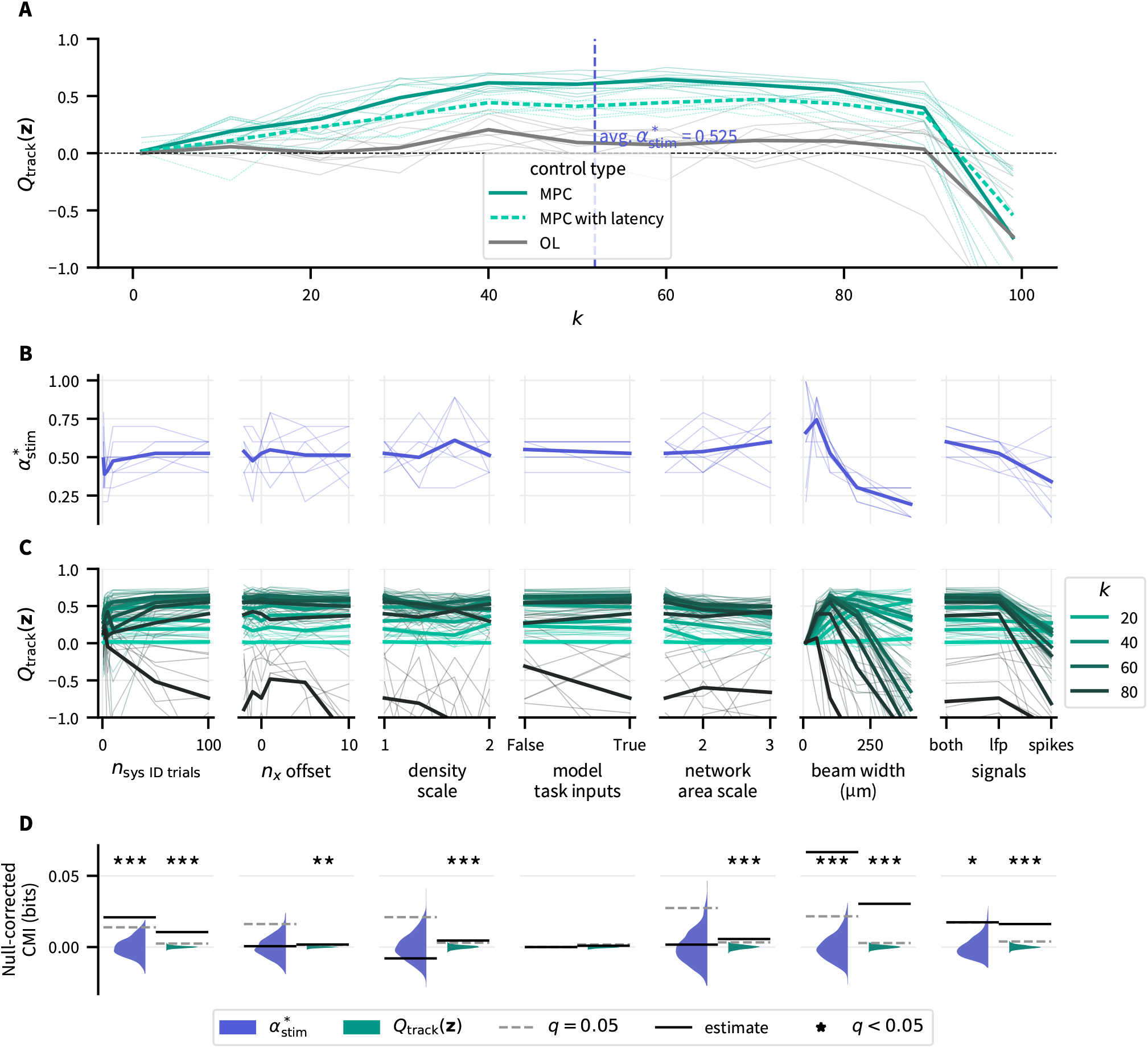
Across a wide range of conditions, latent dynamics were controlled best when stimulation channels were roughly even or slightly higher than recording channels. (A) Performance/*k* curves for the reference configuration show highest performance for a broad peak where *k*≈ *m* or *k* >*m*. Notably, mean performance in the low-*k* regime (α_stim_ <0.3) remained above baseline but below half of peak performance. Performance for the controller with latency was lower for high *k* because the control algorithm punishes *k* much more than *m*. Open-loop (OL) control performed near baseline (peaking around *Q*≈ 0.2) and far below closed-loop control, suggesting it has limited, if any, utility for latent factor control. Here and throughout, thin lines indicate individual seeds and thick lines indicate the mean across seeds. (B) The optimal stimulation/recording balance was relatively consistent with changing exploratory parameters, but decreased particularly for high beam width and spike recording. (C) Control performance was sensitive to all exploratory parameters except task input modeling, depending on α_stim_. (D) Permutation tests, where metric values are shuffled with respect to the parameter values per seed (and channel count for control performance) show statistically significant non-independent relationships. Histograms show CMI estimates for each of 1,000 permutations, solid lines show the point estimate of the unshuffled data, and dashed lines show the *q*= 0.05 significance threshold. *, **, *** indicate *q* <0.05, 0.01, 0.001, respectively.

As with stimulation dimensionality experiments, the main finding (a balanced optimal channel allocation) held across exploratory parameter sweeps (Fig. 5B–D), with significant shifts observed for three parameters:

- *n*_sys ID trials_: More training data shifted 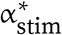 upward, providing the model accuracy needed to leverage higher stimulation channel counts.
- Beam width: Wider illumination beams shifted 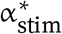 toward recording, perhaps because lower spatial specificity diminished the marginal value of additional stimulation channels.
- Recording signals: Spike-only recordings shifted 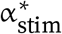 toward recording, likely because sparse neuronal distribution rendered them less informative than LFP. While unusual, LFP has been shown to outperform spikes when analyzing certain experimental datasets [44].

Raw control performance *Q*_track_ showed statistically significant CMI dependence on all exploratory parameters except task input modeling (Fig. 5C,D, Fig. S2)—including the same three parameters shifting 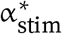 above. For detailed CMI statistics, see Table 2.

**TABLE 2.**
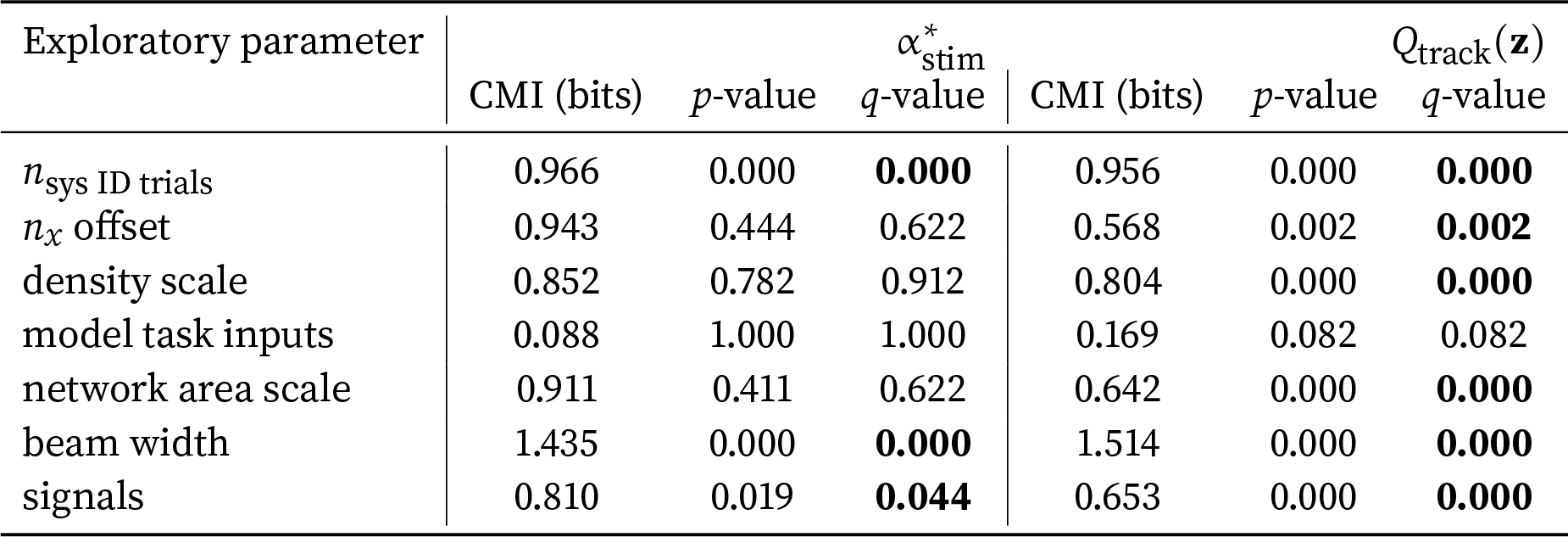
Statistical analysis results for stimulation–recording balance experiments, relating 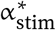 and *Q*_track_(**z**) to exploratory parameters.

| Exploratory parameter | α<sup>*</sup><sub>stim</sub> |  |  | *Q*<sub>track</sub>(**z**) |  |  |
|---|---|---|---|---|---|---|
|  | CMI (bits) | *p*-value | *q*-value | CMI (bits) | *p*-value | *q*-value |
| *n*<sub>sys ID trials</sub> | 0.966 | 0.000 | **0.000** | 0.956 | 0.000 | **0.000** |
| *n*<sub>*x*</sub> offset | 0.943 | 0.444 | 0.622 | 0.568 | 0.002 | **0.002** |
| density scale | 0.852 | 0.782 | 0.912 | 0.804 | 0.000 | **0.000** |
| model task inputs | 0.088 | 1.000 | 1.000 | 0.169 | 0.082 | 0.082 |
| network area scale | 0.911 | 0.411 | 0.622 | 0.642 | 0.000 | **0.000** |
| beam width | 1.435 | 0.000 | **0.000** | 1.514 | 0.000 | **0.000** |
| signals | 0.810 | 0.019 | **0.044** | 0.653 | 0.000 | **0.000** |

## 4. Discussion

In this study we simulated end-to-end CLOC experiments manipulating the latent dynamics of task-trained spiking neural networks, finding several insights to help inform future *in vivo* experiments. First, in stimulation dimensionality experiments, we found that latent factors were controlled effectively with relatively few stimulation channels. Although this confirmed our hypothesis, it challenges prevailing assumptions in systems neuroscience—reflected in heavy investment in single-cell two-photon optogenetics over the mesoscale light array approach used here—that near-per-neuron actuation is required to steer population dynamics. Analogous to how few recording channels capture the dominant variance in population activity, few stimulation channels achieve the majority of potential control. For comparison, in one macaque motor cortex reach dataset, kinematic decoding *R*^2^ exhibited an empirical knee point at ~100 neurons [45], whereas our reference configuration yielded a stimulation knee point of *k*_knee_ = 200. Whether stimulation and recording follow similar scaling laws with respect to task complexity remains an open question [46]. Importantly, although *k*_knee_ independently controlled light-source groups were fewer than the neuron count (~1,024), their number remained larger than the latent dimensionality, highlighting the value of continued development of high-channel-count stimulation technologies.

Second, in stimulation–recording balance experiments, an approximately equal balance of stimulation and recording channels optimized behavioral control under our fixed-total-channel constraint. While predictable from control theory, this balance contrasts sharply with current experimental practice, where multi-contact electrode shanks are routinely paired with a single optical fiber. In our reference simulations, this low-*k* regime (α_stim_< 0.3) yielded substantially less tracking improvement than balanced allocations. The practical lesson is to provision sufficient actuation alongside sensing, not to prescribe a one-to-one hardware tradeoff: modality-specific constraints could instead be represented by *c*_stim_*k*+ *c*_rec_*m* =*B*, with channel costs defined by the limiting resource. Because unequal costs were not tested here, the optimal hardware-specific allocation and its sensitivity to these costs remain open questions. The consistency of these two main findings across parameter sweeps motivates testing their qualitative predictions in *in vivo* preparations, while quantitative channel requirements may differ.

Third, linear model predictive control was able to achieve robust tracking at or near optimal channel allocations (*Q*_track_> 0.60) with little hyperparameter tuning. While more complex behavioral tasks will likely require nonlinear models, these results demonstrate that standard linear methods provide an effective baseline. This noted, we expect varying such parameters as task complexity would strongly affect control performance, though exploring task space was outside the scope of this study. And while compute latency penalized high-channel-count performance in our simulations, this penalty could be mitigated through compiled solvers [47], explicit MPC [48], or neural network feedforward controllers.

Finally, open-loop control performed substantially worse than closed-loop control, offering limited gains at low stimulation channel counts and falling below baseline at high counts. This limited efficacy is notable because our per-trial, long-horizon MPC approach was designed as a strong version of open-loop control. Inter-trial optimization in this manner is even termed closed-loop control in other studies [15, 17, 49], highlighting the power of *intra-trial* feedback over the inter-trial varieties of closed-loop stimulation more commonly reported.

These results should be interpreted in light of several modeling limitations. First, incorporating additional biological features—such as sparse and local connectivity, layer structure, cell-type diversity, multi-compartment morphologies, opsin dynamics, and non-linear behavioral readouts—will be important for tailoring interfaces to specific cortical structures. Second, our framework assumes rate-coding dynamics; neural circuits relying significantly on spike-timing codes will likely require alternative modeling and control formulations. Finally, extending simulations to tasks with varying cognitive demands and exploring two-photon holographic stimulation [24, 50–53] will further test generalizability. Beyond addressing these limitations, however, there are a number of exciting future directions for control of latent dynamics. First, causal inference is needed to achieve the ultimate scientific goal of controlling latent factors—to assess whether they in fact drive correlated behavior or neural activity. In contrast to the simple scenario tested here, where the latent state *X* completely caused behavior *Z*, causality in real neural systems can only be characterized with rigorous statistical methods. Such causal inference methods have only recently begun to be applied to systems neuroscience data [54, 55]. Another promising direction, alluded to in Sec. 1, consists in constraining perturbations to remain on the full neural manifold [18, 19]. This might involve quantifying distance from the manifold—in tractable fashion amenable to real-time optimal control—and modeling techniques tailored to manifold structure [56, 57]. Besides off-manifold activity, tissue damage through heat, ionic imbalance, etc. could be another useful objective for the controller to minimize.

Finally, our findings establish three concrete predictions for *in vivo* validation:

a. Low-dimensional stimulation achieves the vast majority of potential latent control.
b. Balancing stimulation and recording channels optimizes behavioral tracking.
c. Linear model predictive control with ≥100 total (stimulation plus recording) channels can reduce trajectory tracking error by ~50% when evoking stereotypical motor behaviors. (Though this prediction is much more speculative given the limitations listed above.)

If the approach described here proves feasible *in vivo*, the latent dynamics modeling field could enter a new phase of causal characterization. More generally, leveraging goal-driven latent dynamics modeling [34, 44, 58, 59] to discover neural features, together with optimal control to causally characterize those candidate features, could provide a foundation for high-throughput, data-driven neuroscience. Such a scenario could represent a “bitter lesson” [60] moment for neuroscience, where scaling up sensing (electrophysiology, two-photon imaging, etc.), actuation (optogenetics [4, 5]), and rich behavioral data [61–71] fuels progress outpacing manual efforts, at least for hypotheses of the form “X population does Y”. Regardless, combining machine learning and control theory with multi-input/-output tools represents a vital opportunity to unlock deeper understanding of the systems and circuits of the brain.

## Acknowledgments

This work was supported by National Institutes of Health (NIH) NINDS CRCNS grant R01NS115327. K.A.J. was additionally supported by the Georgia Tech/Emory NIH/NIBIB Training Program in Computational Neural Engineering (T32EB025816).

## Competing interests

C.J.R. is a consultant to Motif Neurotech, Inc. C.P. is a Senior Adviser at Inductive Technologies. Inductive Technologies was not involved in this study.

## A. Supplement

### A.1 Cleo interface details

In stimulation dimensionality experiments, to partition the grid of light sources into clusters as evenly as contiguously as possible, we used constrained Voronoi partitioning. Each cluster had a fixed number of channels and channels were assigned to slots by minimizing the total squared Euclidean distance between neurons and cluster centroids using SciPy’s `linear_sum_assignment()` [72] function and repeating until convergence. Cluster sizes were determined using integer division, i.e., *n*_cluster_= ⌊1024/*k*⌋, and the remaining 1024− ⌊1024 /*k* ⌋channels were assigned one each to the first so many clusters.

In stimulation–recording balance experiments, on the other hand, we needed members of each group (two groups, light sources and electrodes) to be as far apart from each other as possible so each modality could best cover the available area. We initially assigned distributed *m* electrode and *k* light source randomly, then iteratively picked a random electrode/light pair to consider swapping if it reduced the objective of minimizing “force” between members of the same group, as between same-charged particles

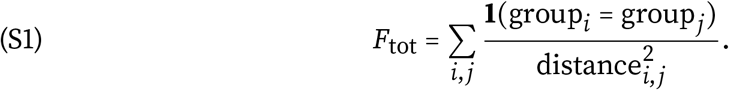

The algorithm continued until 100 successive pair samples that did not improve the objective function, i.e., were not swapped.

### A.2. Kernel density estimation

KDE was performed for heterogeneous data types by mixing kernels: the Wang-Ryzin kernel [73] for ordered discrete variables (most exploratory parameters), the Aitchison-Aitken kernel [74] for unordered discrete variables (random seed, electrophysiology signals, and task input modeling), and a Gaussian kernel for continuous variables (control performance and other metrics), using the implementation of the `statsmodels` Python package [75]. Various heuristics were used to select bandwidths. For the Aitchison-Aitken kernel, we set 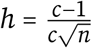 where *c* is the number of categories and *n* is the number of samples. For the Wang-Ryzin kernel, we set 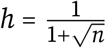. For the Gaussian kernel, we used Silverman’s rule of thumb [76]. ^+^

### A.3. Supplemental table

**TABLE S1.**
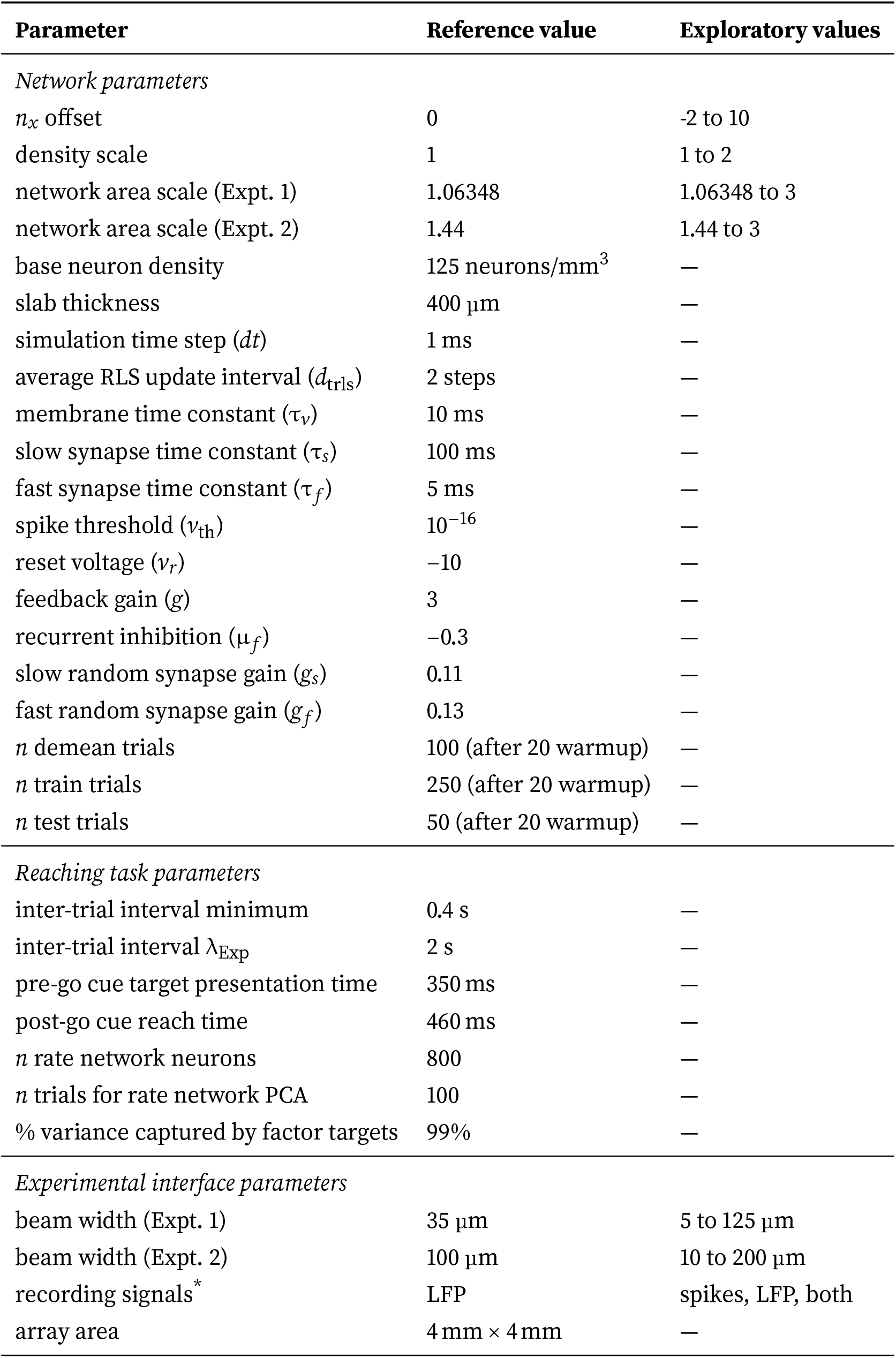
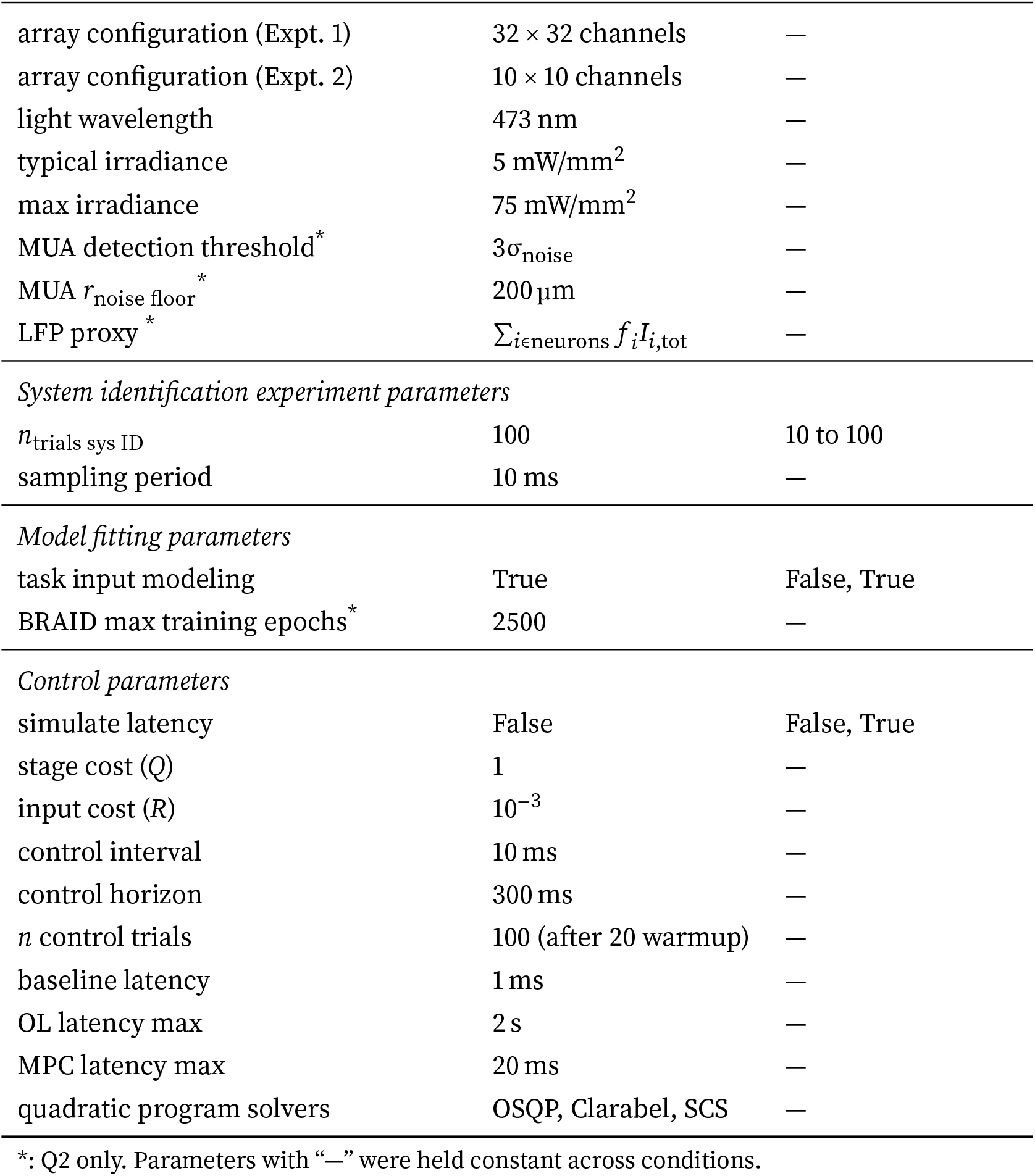
Model and experiment parameters.

| Parameter | Reference value | Exploratory values |
|---|---|---|
| *Network parameters* |  |  |
| *n*<sub>*x*</sub> offset | 0 | -2 to 10 |
| density scale | 1 | 1 to 2 |
| network area scale (Expt. 1) | 1.06348 | 1.06348 to 3 |
| network area scale (Expt. 2) | 1.44 | 1.44 to 3 |
| base neuron density | 125 neurons/mm<sup>3</sup> | — |
| slab thickness | 400 µm | — |
| simulation time step (*dt*) | 1 ms | — |
| average RLS update interval (*d*<sub>trls</sub>) | 2 steps | — |
| membrane time constant (τ<sub>*v*</sub>) | 10 ms | — |
| slow synapse time constant (τ<sub>*s*</sub>) | 100 ms | — |
| fast synapse time constant (τ<sub>*f*</sub>) | 5 ms | — |
| spike threshold (*v*<sub>th</sub>) | 10<sup>−16</sup> | — |
| reset voltage (*v*<sub>*r*</sub>) | −10 | — |
| feedback gain (*g*) | 3 | — |
| recurrent inhibition (μ<sub>*f*</sub>) | −0.3 | — |
| slow random synapse gain (*g*<sub>*s*</sub>) | 0.11 | — |
| fast random synapse gain (*g*<sub>*f*</sub>) | 0.13 | — |
| *n* demean trials | 100 (after 20 warmup) | — |
| *n* train trials | 250 (after 20 warmup) | — |
| *n* test trials | 50 (after 20 warmup) | — |
| *Reaching task parameters* |  |  |
| inter-trial interval minimum | 0.4 s | — |
| inter-trial interval λ<sub>Exp</sub> | 2 s | — |
| pre-go cue target presentation time | 350 ms | — |
| post-go cue reach time | 460 ms | — |
| *n* rate network neurons | 800 | — |
| *n* trials for rate network PCA | 100 | — |
| % variance captured by factor targets | 99% | — |
| *Experimental interface parameters* |  |  |
| beam width (Expt. 1) | 35 µm | 5 to 125 µm |
| beam width (Expt. 2) | 100 µm | 10 to 200 µm |
| recording signals<sup>*</sup> | LFP | spikes, LFP, both |
| array area | 4 mm × 4 mm | — |
| array configuration (Expt. 1) | 32 × 32 channels | — |
| array configuration (Expt. 2) | 10 × 10 channels | — |
| light wavelength | 473 nm | — |
| typical irradiance | 5 mW/mm<sup>2</sup> | — |
| max irradiance | 75 mW/mm<sup>2</sup> | — |
| MUA detection threshold<sup>*</sup> | 3σ<sub>noise</sub> | — |
| MUA *r*<sub>noise floor</sub><sup>\*</sup> | 200 µm | — |
| LFP proxy<sup>*</sup> | ∑<sub>*i*∈neurons</sub> *f*<sub>*i*</sub>*I*<sub>*i*,tot</sub> | — |
| *System identification experiment parameters* |  |  |
| *n*<sub>trials sys ID</sub> | 100 | 10 to 100 |
| sampling period | 10 ms | — |
| *Model fitting parameters* |  |  |
| task input modeling | True | False, True |
| BRAID max training epochs<sup>*</sup> | 2500 | — |
| *Control parameters* |  |  |
| simulate latency | False | False, True |
| stage cost (*Q*) | 1 | — |
| input cost (*R*) | 10<sup>−3</sup> | — |
| control interval | 10 ms | — |
| control horizon | 300 ms | — |
| *n* control trials | 100 (after 20 warmup) | — |
| baseline latency | 1 ms | — |
| OL latency max | 2 s | — |
| MPC latency max | 20 ms | — |
| quadratic program solvers | OSQP, Clarabel, SCS | — |

\*: Q2 only. Parameters with “—” were held constant across conditions.

### A.4. Supplemental figures

**FIGURE S1.**
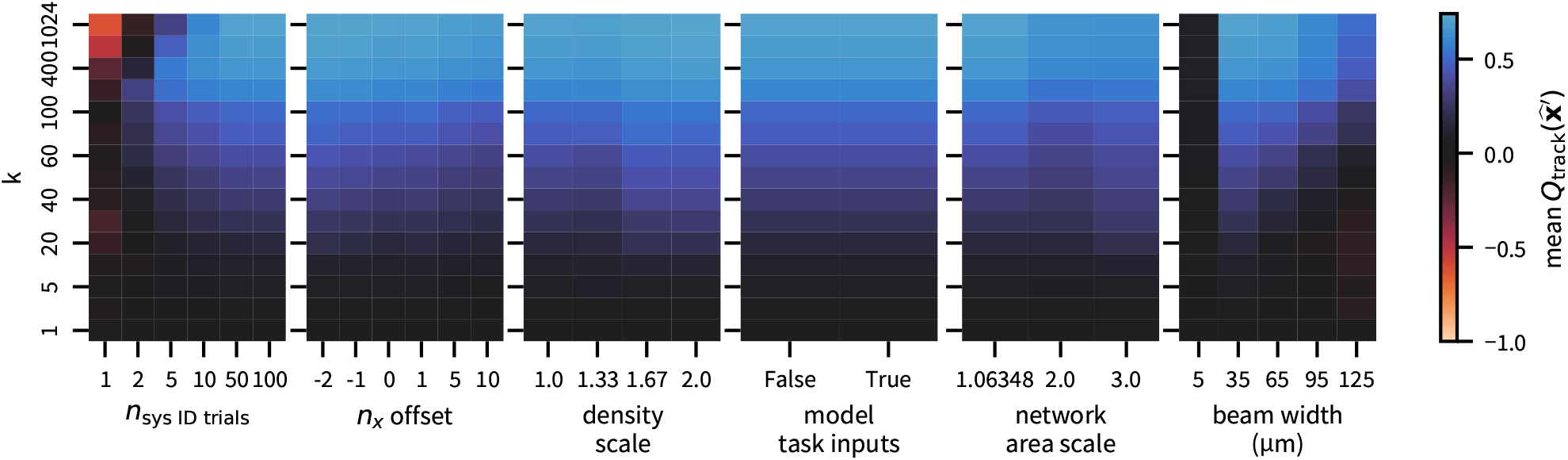
Control performance heatmaps for stimulation dimensionality experiments. Each plot shows the results for the reference condition averaged over seeds, varying only the exploratory parameter (columns) and number of stimulation channels *k* (rows). Extreme values outside the color bar range are displayed directly on the heat map.

**FIGURE S2.**
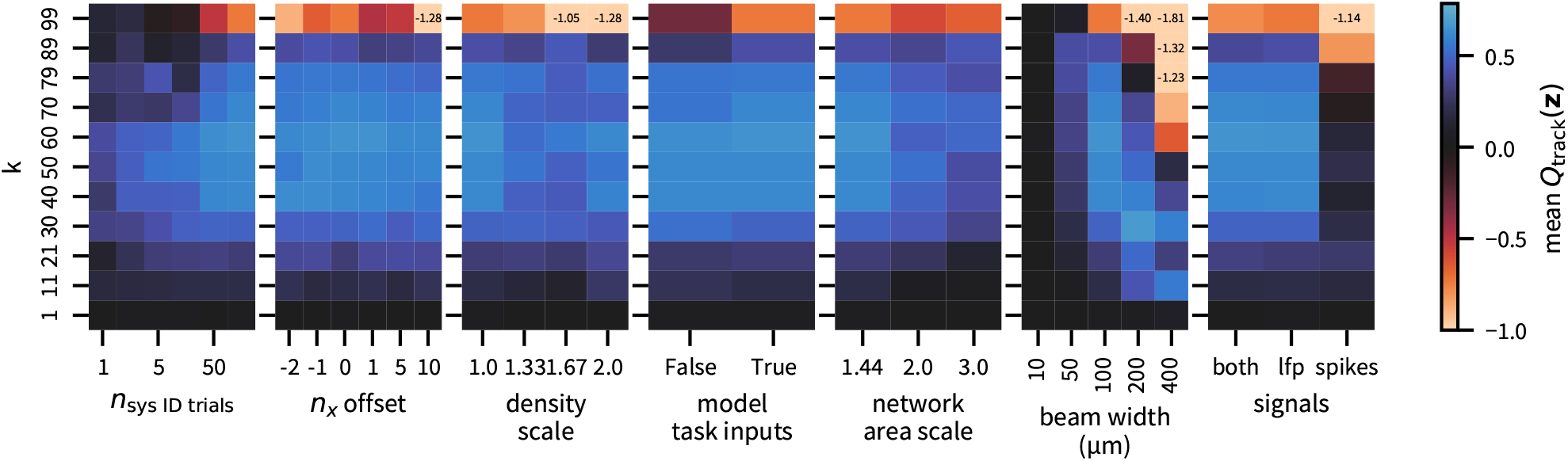
Control performance heatmaps for stimulation–recording balance experiments. Each plot shows the results for the reference condition averaged over seeds, varying only the exploratory parameter (columns) and number of stimulation channels *k* (rows). Extreme values outside the color bar range are displayed directly on the heat map.

## Footnotes

1 Here and throughout the paper, metrics are reported as mean ±SEM.

2 Reference configuration parameters are defined in Table S1; see Sec. 2.7.

3 Reach accuracy evaluated via nearest-neighbor trajectory classification; see Sec. 2.8

## Notes

https://github.com/siplab-gt/control-of-latent-dynamics

